# *De novo* transcriptome characterization of a sterilizing trematode parasite (*Microphallus* sp.) from two species of New Zealand snails

**DOI:** 10.1101/086801

**Authors:** Laura Bankers, Maurine Neiman

## Abstract

Snail-borne trematodes represent a large, diverse, and evolutionarily, ecologically, and medically important group of parasites, often imposing strong selection on their hosts and causing host morbidity and mortality. Even so, there are very few genomic and transcriptomic resources available for this important animal group. We help to fill this gap by providing transcriptome resources from trematode metacercariae infecting two congeneric snail species, *Potamopyrgus antipodarum* and *P*. *estuarinus*. This genus of New Zealand snails has gained prominence in large part through the development of P. antipodarum and its sterilizing trematode parasite *Microphallus livelyi* into a textbook model for host-parasite coevolutionary interactions in nature. By contrast, the interactions between *Microphallus* trematodes and *P. estuarinus*, an estuary-inhabiting species closely related to the freshwater *P*. antipodarum, are relatively unstudied. Here, we provide the first annotated transcriptome assemblies from *Microphallus* isolated from *P*. *antipodarum* and *P*. *estuarinus*. We also use these transcriptomes to produce genomic resources that will be broadly useful to those interested in host-parasite coevolution, local adaption, and molecular evolution and phylogenetics of this and other snail-trematode systems. Analyses of the two *Microphallus* transcriptomes revealed that the two trematode isolates are more genetically differentiated from one another than are *M. livelyi* infecting different populations of *P. antipodarum*, suggesting that the *Microphallus* infecting *P*. *estuarinus* represent a distinct lineage. We also provide a promising set of candidate genes likely involved in parasitic infection and response to salinity stress.

## INTRODUCTION

Snail-borne trematodes are important sources of morbidity and mortality in animals, including humans (Hotez 2013; Jurberg and Brindley 2015). For example, infections by trematodes in the genus *Schistosoma* affect more than 200 million people in sub-Saharan Africa (Rollinson *et al.* 2013). Trematodes and their mollusk intermediate hosts are also important models for the study of host-parasite coevolutionary interactions (*e.g.*, Lively and Jokela 1996; King *et al*. 2009). While digenetic trematodes (parasitic flatworms) are one of the largest groups of metazoan parasites, with ∼18,000 species (Cribb *et al*. 2001), the genomic and transcriptomic resources needed to understand trematode evolution, host-parasite coevolution, and the genomic basis of trematode infection are only available for ten of these species (NCBI, October, 2016). Further development of such resources for this medically and scientifically important group of parasites is a necessary component of characterizing the molecular and genomic underpinnings of parasitic infection and host-parasite coevolutionary interactions.

New Zealand snails in the genus *Potamopyrgus* are often the intermediate host of sterilizing trematode parasites in the genus *Microphallus* (Winterbourn 1973). Here, we focus on *Microphallus* ‘*livelyi*’ (Hechinger 2012), a species of digenetic trematode that uses *Potamopyrgus antipodarum* as an intermediate host and dabbling ducks (Anatinae) as the final host (Dybdahl and Lively 1996; Lively and Jokela 1996; King *et al*. 2009; Osnas and Lively 2011). Dabbling ducks are infected by *M. livelyi* while foraging, typically consuming infected *P. antipodarum* from submerged rocks and aquatic vegetation. The trematodes lay sexually produced eggs within the ducks, which are excreted with duck feces (Osnas and Lively 2011). The next round of snail infection begins after detritus-feeding *P. antipodarum* consume *M. livelyi* eggs while foraging along lake bottoms and aquatic vegetation (Lively and Jokela 1996). Following snail consumption, the *M. livelyi* eggs clonally develop into thousands of encysted larvae (metacercariae) that rapidly sterilize the snail (Winterbourn 1973; King *et al.* 2009). Because *M. livelyi* that cannot infect the first snail by which they are ingested die before completing their lifecycle, and because successful infection means that the *M. livelyi* metacercariae sterilize the snail, there is strong reciprocal antagonistic selection pressures between these two species (King *et al.* 2011). The metacercariae life stage thus represents a crucial time point for *Microphallus* both from the perspective of its life cycle and with respect to its coevolutionary interactions with *P. antipodarum.*

Dabbling ducks carry *M. livelyi* as they migrate between New Zealand lakes, thereby increasing the level of parasite gene flow relative to that of *P. antipodarum* (Dybdahl and Lively 1996). Accordingly, there is very little population structure in *M. livelyi* relative to *P. antipodarum* (Dybdahl and Lively 1996), which itself features marked among-lake genetic structure (Paczesniak *et al*. 2013). *Microphallus livelyi* is locally adapted to sympatric *P. antipodarum* both among and within lake populations (Lively *et al*. 2004; King *et al*. 2009). Within lake populations, shallow regions tend to show both relatively high infection frequency as well as pronounced local adaptation (King *et al*. 2009). Consistent with the expectation of local adaptation, *M. livelyi* infection is associated with population-specific patterns of gene expression and genetic differentiation in *P. antipodarum* (Bankers *et al*. in review).

How interactions between *M. livelyi* and *P. antipodarum* play out in *Microphallus* are not well characterized. Some initial insight into the mechanisms underlying *Microphallus* infectivity come from the several studies suggesting that coevolutionary interactions between *P. antipodarum* and *M. livelyi* fit a “matching alleles” infection genetics model, whereby there are no universally infective parasites or resistant hosts (Dybdahl and Krist 2004; Lively *et al*. 2004; King et al. 2009). More specifically, *M. livelyi* infection success is thought to be determined by whether the *M. livelyi* genotype at the loci involved in mediating infection match the corresponding loci in *P. antipodarum,* enabling successful *M. livelyi* to evade detection by *P. antipodarum* defenses. While matching alleles mechanisms for infection have been demonstrated in several host-parasite interactions, whether these mechanisms are mediated by gene expression and/or by sequence evolution remains unclear in *P. antipodarum* or in any other system (Carmona *et al*. 2015; Routtu and Ebert 2015).

The transcriptomic resources described here, the first genomic resources for the *Microphallus* genus of which we are aware, will provide a valuable starting point to characterize the underpinnings of *M. livelyi* infection and the coevolutionary interactions between *M. livelyi* and *P. antipodarum*. We present annotated transcriptomes assemblies from *M. livelyi* metacercariae isolated from *P. antipodarum* as well as from *Microphallus* of unknown species isolated from *P. estuarinus*, a closely related species native to New Zealand estuaries. While the coevolutionary interactions between *Microphallus livelyi* and *P. antipodarum* are well characterized, much less is known about the *Microphallus* that infect *P. estuarinus* (Winterbourn 1973; Hechinger 2012). Expanding the genomic resources available for these two closely related types of *Microphallus* will allow for the characterization of genes that contribute broadly to infection processes as well as genes that play species-specific roles in host-parasite interactions.

We identified and annotated 7584 pairs of one-to-one orthologs, analyzed levels of genetic variation within and genetic differentiation between the two types of *Microphallus* at these genes, and performed gene ontology-based functional comparisons between the two isolates. We also provide a broad set of genomic resources that set the stage for future studies aimed at characterizing the molecular underpinnings of the coevolutionary interactions between *Microphallus* and *Potamopyrgus*.

## MATERIALS AND METHODS

### Sample collection and RNA sequencing

We isolated *M. livelyi* from four individual adult female *Potamopyrgus antipodarum* (“PA-*Microphallus*”) collected in January of 2013 in shallow regions (<1m lake depth) of each of the two New Zealand lakes Alexandrina and Kaniere, for a total of eight PA-*Microphallus* isolates. We chose these lakes because their *P. antipodarum* populations contain relatively high frequencies of *M. livelyi* infection (10-20%; Vergara *et al*. 2013). We also isolated *Microphallus* from a single adult female *Potamopyrgus estuarinus* (“PE-*Microphallus*”) collected from the Ashley River estuary (near Christchurch, New Zealand) in January of 2015.

We dissected individual *Potamopyrgus* snails to determine infection status, and saved parasite tissue from snails containing stage 5 *Microphallus* infections (fully formed metacercariae that fill the entire snail body cavity; http://www.indiana.edu/%7Ecurtweb/trematodes/DATA_KEY.HTM). We separated parasite from snail tissue with a forceps and used a micropipette to isolate metacercariae cysts. We immediately placed the metacercariae in RNAlater® Solution (Life Technologies Corporation) at 4°C for 24 hours and subsequently stored the RNAlater®-submerged metacercariae at - 80°C (according to manufacturer protocol) until RNA extraction. The eight PA-*Microphallus* metacercariae isolates were pooled in one tube. We had to use this pooling strategy for the PA-*Microphallus* because at this time (early 2013), pooling of cysts from multiple snails was necessary to obtain a sufficient amount of RNA to perform RNA sequencing. Technological advances in library preparation and RNA sequencing between 2013 and 2015 meant that we were able to obtain a sufficient amount of RNA from the metacercariae isolated from a single *P. estuarinus* individual for RNA sequencing.

We extracted RNA from metacercariae following the TRIzol protocol (Chomczynski and Sacchi 1987; Chomczynski 1993). We assessed RNA quantity and quality using a Bio-Rad Experion Automated Electrophoresis Station and Experion RNA analysis kit, requiring an RQI minimum of 2 µg of total RNA (following manufacturer protocol). We prepared cDNA libraries following the Illumina Truseq LS protocol (Illumina, San Diego, CA, 2012) for the PA-*Microphallus* and PE-*Microphallus* samples. We barcoded the PA-*Microphallus* library and pooled the barcoded library with 11 other barcoded samples that were part of a different project in one Illumina HiSeq 2000 lane in 2013. For the PE-*Microphallus* library, we barcoded the library and pooled the barcoded library with five other samples that were part of a different project in one Illumina HiSeq 2000 lane in 2015. We performed 2x100 bp paired-end RNA sequencing for both *Microphallus* libraries (Illumina, San Diego, CA, 2012). We obtained 31635450 paired-end reads (mean read length = 101 bp) for PA-*Microphallus* and 41615863 paired-end reads for PE-*Microphallus* (mean read length = 126 bp).

### Transcriptome assembly and annotation

We used FASTQC (Andrews 2010) to assess RNA sequencing quality and the FASTX Toolkit (Gordon and Hannon 2010) to trim adapter sequences and remove poor-quality reads from raw RNA-Seq data, requiring a mean Phred score ≥ 20. After quality filtering, 31096527 paired-end reads remained for PA-*Microphallus* and 40413790 paired-end reads remained for PE-*Microphallus*. We used these filtered reads to generate *de novo* transcriptome assemblies for each of the two *Microphallus* isolates.

We performed the Trinity *in silico* normalization, requiring a minimum Kmer coverage of 2.0x and maximum Kmer coverage of 10x (following Haas *et al*. 2013), and otherwise default parameters (Gabherr *et al*. 2011). We then generated preliminary *de novo* transcriptome assemblies for each isolate using Trinity software with default parameters (Haas *et al*. 2013). Preliminary assemblies contained 65665 contigs and 226896 contigs for PA-*Microphallus* and PE-*Microphallus*, respectively. We used the Trinity plugin TransDecoder with default parameters to annotate likely protein-coding regions, identify long ORFs of putative genes, and filter miscalled isoforms (Haas *et al*. 2013). Next, we used hierarchical clustering based on sequence identity to further reduce redundancy in the transcriptome assemblies, as implemented by CD-HIT-EST (Huang *et al*. 2010). To reduce the likelihood of collapsing truly non-redundant transcripts, we used a sequence identity of 0.95 and word size of eight nucleotides (*e.g*., Li *et al*. 2002). These steps resulted in assemblies containing 15435 contigs for PA-*Microphallus* and 29564 contigs for PE-*Microphallus*. We mapped reads back to their respective filtered transcriptome assemblies using Tophat2 (Trapnell *et al.* 2013) and used Samtools depth (Li *et al.* 2012) to estimate mean per-base coverage (+/− SD) for sites covered by at least 1x. Mean coverage for the transcriptome assemblies were 101.4x (+/− 330.1) for PA-*Microphallus* and 147.5x (+/− 437.6) for PE-*Microphallus*. The nearly two-fold higher number of transcripts for PE-*Microphallus* relative to PA-*Microphallus* was likely a combination of advances in sequencing chemistry and technology between 2013 and 2015 as well as higher sequencing depth for PE-*Microphallus.* We annotated the transcriptomes using blastx with an E-value cutoff of 1e-6, followed by Blast2GO (using default parameters) to assign GO terms to blastx-annotated transcripts (Conesa *et al*. 2005). For PA-*Microphallus*, we obtained both blastx and GO annotations for 9272 transcripts and only blastx annotations for an additional 3814 transcripts. For PE-*Microphallus* we obtained both blastx and GO annotations for 13778 transcripts and only blastx annotations for an additional 8822 transcripts. This analysis revealed 143 transcripts involved in photosynthesis in the PE-*Microphallus* assembly (none in PA-*Microphallus*), representing potential contaminants in the snail gut from which the PE-*Microphallus* was isolated. These 143 transcripts were filtered out of subsequent analyses. Finally, we used BUSCO (Simão *et al*. 2015) to assess the completeness of our transcriptome assemblies. This program uses tblastn to identify conserved single-copy orthologs among groups of organisms. We used the BUSCO OrthoDB Metazoan database to search our assemblies for the presence, absence, duplication, and/or fragmentation of 843 conserved single-copy Metazoan orthologs.

### Identification of orthologous transcripts

We used “reciprocalblast_allsteps.py” (Warren *et al.* 2014) to identify putative orthologous transcripts between the two *Microphallus* transcriptomes based on one-to-one reciprocal best blastn hits. We performed reciprocal blastn searches between the two transcriptomes with an E-value cutoff of 1e-20 and identified pairs of transcripts that were reciprocally one another’s top blastn hit and that occurred once and only once in each transcriptome query. We identified 11160 of such transcript pairs, which we will hereafter refer to as “one-to-one orthologs.” Using a custom python script (github.com/jsharbrough/grabContigs), we extracted the 11060 orthologous sequences from each *Microphallus* transcriptome to generate putative ortholog transcriptomes for each isolate. To improve the accuracy of our ortholog predictions, we then used blastx against the NCBI nr database with an E-value cutoff of 1e-6 (Camacho *et al*. 2009) to annotate the two transcriptomes and filter out putative orthologs that were annotated as different genes. This step resulted in 7935 remaining putative one-to-one orthologs. To further ensure that we accurately predicted orthologs, we also performed reciprocal shortest molecular distance comparisons between the 7935 pairs of transcripts using “reciprocal_smallest_distance.py” (Wall *et al.* 2003). Here, we inferred orthologs using a combination of global sequence alignments between the two transcriptomes and maximum likelihood-based evolutionary pairwise distances, and identified orthologous pairs of transcripts as those sets of transcripts with the shortest pairwise molecular distance between them relative to their pairwise distances from any of the other 7934 transcripts in the dataset (Wall *et al.* 2003). This filtering step left 7584 remaining pairs of one-to-one orthologous transcripts that met all three of our criteria: (1) reciprocal best blastn hits, (2) the same blastx annotation, and (3) shared the shortest pairwise evolutionary distance relative to all other transcripts. There were 7851 and 21980 transcripts for PA-*Microphallus* and PE-*Microphallus*, respectively, that did not meet these three criteria (hereafter, “non-one-to-one orthologs”). We extracted the 7584 orthologous transcripts from each reference transcriptome to generate ortholog transcriptomes for each of the two *Microphallus* datasets.

### Comparisons of genetic variation and genetic differentiation

To assess levels of genetic differentiation and nucleotide heterozygosity (Watterson’s θ) within and between the two isolates, we used Tophat2 (Trapnell *et al.* 2013) to map *Microphallus* reads to the ortholog transcriptomes. Because we had two different ortholog transcriptomes that represented different numbers of *Microphallus* individuals, we performed four mapping combinations: (1) PA-*Microphallus* reads to the PA-*Microphallus* ortholog transcriptome, (2) PE-*Microphallus* reads to the (3) PE-*Microphallus* reads to the PA-*Microphallus* ortholog transcriptome, and (4) PA-*Microphallus* reads to the PE-*Microphallus* ortholog transcriptome. This design allowed us to estimate Watterson’s θ within and between the two types of *Microphallus* while accounting for the ortholog transcriptome to which the reads were mapped. Next, we used Picard Tools addition of read groups and removal of duplicate reads from each bam file. We then used GATK v.1.119 (http://picard.sourceforge.net) to process and filter the bam files. This step included the (McKenna *et al*. 2010; DePristo *et al*. 2011; Van der Auwera *et al*. 2013) to realign reads around indels and to reassign mapping quality. Finally, we used Samtools (Li *et al.* 2012) to sort each bam file by coordinates relative to the ortholog transcriptome to which they were mapped and to generate mpileups.

We generated separate mpileup files for each of the four bam files and used Popoolation (Kofler *et al*. 2011a) for further analyses. Popoolation is a suite of programs designed to handle pooled sequencing data and is able to account for the number of individuals sequenced. First, we used Popoolation’s Variation-at-position.pl script (which is not available in newer program versions) to calculate Watterson’s θ within and between the two *Microphallus* types, using each of the four bam Bonferroni-corrected Mann-Whitney U-tests and Kolmogorov-Smirnov tests, respectively, as files to take into account the ortholog transcriptome to which reads were mapped. We compared the medians and distributions of θ for all six possible pairwise comparisons of the four bam files with (Popoolation2) (Kofler *et al*. 2011b) to generate synchronized mpileups (mpileup2sync.pl) for each implemented in SPSS Statistics v. 23. We then used the most up-to-date version of Popoolation pair of bam files associated with each ortholog transcriptome (*i.e.*, PA-*Microphallus* and PE-*Microphallus* mapped to the PA-*Microphallus* ortholog transcriptome, and PA-*Microphallus* and PE-*Microphallus* mapped to the PE-*Microphallus* ortholog transcriptome). Next, we used the Popoolation2 fst-sliding.pl to calculate *F*_*ST*_ per SNP to assess levels of relative genetic differentiation between the two *Microphallus* types. We used these *F*_*ST*_ values to determine mean *F*_*ST*_ per SNP between PA-*Microphallus* and PE-*Microphallus* relative to both of the ortholog transcriptomes. We applied outlier analyses within SPSS Statistics v. 23 to identify transcripts containing *F*_*ST*_ outlier SNPs, which can be interpreted as representing significant genetic differentiation for these genes between these two types of *Microphallus.* Finally, we used blastx and Blast2GO to annotate *F*_*ST*_ outlier-containing transcripts and determine putative functions.

### GO and KEGG-based functional comparisons of transcriptomes and orthologs

We used blastx (Camacho *et al*. 2009) and Blast2GO (Conesa *et al.* 2005) to annotate the orthologous sequences for both *Microphallus* types. We also annotated the transcripts from each transcriptome that did not fall into orthologous gene sets (“non-one-to-one orthologs”). We then performed GO functional enrichment analyses, using Fisher’s Exact tests as implemented by Blast2GO (Conesa *et al.* 2005), to identify over *vs.* underrepresented functional groups in the set of orthologous transcripts relative to the rest of the transcriptome and for the set of non-one-to-one orthologous transcripts relative to the rest of the transcriptome. All functional enrichment analyses were performed on both *Microphallus* samples. This analysis approach allowed us to compare over *vs.* underrepresented functional groups for orthologous and non-orthologous transcripts between the two types of *Microphallus* in order to identify which functional groups were expressed in both *Microphallus* isolates *vs.* those functional groups that were uniquely enriched in each *Microphallus* type. We also predicted KEGG (Kyoto Encyclopedia of Genes and Genomes database) pathways for each transcriptome as implemented by Blast2GO. This analysis allowed us to (1) identify potential among-gene interactions, (2) identify sets of interacting genes of relevance to the *Microphallus* life cycle stage that we sequenced, and (3), assess whether different pathways are being expressed between the two isolates. Finally, we used Blast+ to generate local BLAST databases for both annotated transcriptomes.

### Microsatellite loci and primer prediction

We used PrimerPro V.1.0 (https://webdocs.cs.ualberta.ca/∼yifeng/primerpro/) to identify potential microsatellite loci in the two *Microphallus* transcriptomes. We limited these analyses to the ortholog We used PrimerPro V.1.0 (https://webdocs.cs.ualberta.ca/∼yifeng/primerpro/) to identify potential transcriptomes to ensure the transcripts in which we identified microsatellites were distinct loci and to allow future direct comparisons among *Microphallus* isolates. The PrimerPro V.1.0 pipeline first predicts microsatellite loci by identifying simple sequence repeats (SSRs) using MISA (MIcroSAtellite Identification Tool; http://pgrc.ipk-gatersleben.de/misa/misa.html) and then uses Primer3 (Untergasser *et al.* 2012) and BLAST (Camacho *et al.* 2009) to generate primers isolating these loci. After these steps, PrimerPro V.1.0 outputs the putative microsatellites, the primer sequences, their transcriptomic locations, and the primer properties. Following Qiu *et al.* (2016), we allowed for a minimum of ten repeats for mononucleotide repeats, six for dinucleotide repeats, five for trinucleotide repeats, four for tetranucleotide repeats, three for pentanucleotide repeats, and three for hexanucleotide repeats, and identified compound microsatellites as instances where there was more than one microsatellite separated by less than or equal 150 nucleotides.

### Data Availability

The raw RNA-Seq reads will be deposited on NCBI SRA. The annotated transcriptome assemblies will be deposited on GenBank. Blast databases will be made available at http://bioweb.biology.uiowa.edu/neiman/Potamomics.php. VCF-formatted SNP data and microsatellite loci with predicted primer details will be deposited on Dryad. GO annotation, enrichment, and KEGG pathway analyses are provided in the online supplementary information. Custom python script is available at: github.com/jsharbrough/grabContigs.

## RESULTS AND DISCUSSION

### Transcriptome assembly and annotation

274

We generated *de novo* transcriptome assemblies for stage 5 *Microphallus* metacercariae isolated from field-collected *Potamopyrgus antipodarum* (PA-*Microphallus*) and *Potamopyrgus estuarinus* (PE-*Microphallus*). Our final transcriptome assemblies contained 15435 and 29564 transcripts for PA-*Microphallus* and PE-*Microphallus*, respectively. The assembly for PE-*Microphallus* contained ∼2x as many transcripts and was 33% longer than the assembly for PA-*Microphallus*. The PA-*Microphallus* assembly had, on average, 22% longer transcripts and 23% larger N50 than the PE-*Microphallus* assembly (Table 1). The marked differences in these assembly statistics are at least in part linked to differences in sequencing depth and technological advances between the two rounds of sequencing but could also reflect, for example, biological differences between *Microphallus* found in freshwater (inhabiting *P. antipodarum) vs.* estuarine (inhabiting *P. estuarinus*) habitats.

We obtained both blastx and GO annotations for 60.1% and 46.6% of transcripts and only blastx annotations (including those with GO mapping, but not GO annotation) for an additional 24.6% and 29.8% of transcripts for PA-*Microphallus* and PE-*Microphallus*, respectively. There was a mean of ∼3.4 (SD) GO annotations per transcripts per transcriptome. The remaining 15.2% of PA-*Microphallus* transcripts and 23.6% of PE-*Microphallus* transcripts were unable to be annotated, which is typical for *de novo* assemblies of non-model taxa (*e.g.*, Guo *et al.* 2015; Theissinger *et al*. 2016). The species that were most frequently the top blast hit for a given transcript were most often other trematodes (10053 transcripts, 77.4% for PA-*Microphallus*; 12342 transcripts, 54.9% for PE-*Microphallus)*, or non-trematode members of Platyhelminthes (an additional 332 transcripts, 2.6% for PA-*Microphallus*; 382 transcripts, 1.7% for PE-*Microphallus*) (Table S1).

We used BUSCO to estimate the completeness of our transcriptome assemblies based on the presence, absence, duplication, and/or fragmentation of conserved Metazoan genes. Both transcriptomes contain complete transcripts for ∼68% of the 843 conserved single-copy Metazoan genes in the BUSCO database. Our transcriptome assemblies contain gene fragments for an additional 7.8% (PA-*Microphallus*) and 14.5% (PE-*Microphallus*) of the conserved single-copy Metazoan genes, for a total of 76.5% (PA-*Microphallus*) and 82.4% (PE-*Microphallus*) of the 843 BUSCO genes represented in the transcriptomes (Table 2). Considered together with our assembly statistics (Table 1), these results indicate that our transcriptomes are of good quality, are reasonably complete, and are qualitatively similar to other recently published *de novo* transcriptome assemblies (*e.g.*, Guo *et al.* 2015; Hara *et al.* 2015; Tassone *et al.* 2016; Theissinger *et al.* 2016), despite the differences in transcript number, coverage, and N50 between the two transcriptome assemblies. Because we analyzed transcriptomes generated from only one stage of infection from a parasite with a complex life cycle, at least some of the missing BUSCO genes likely represent genes that are not expressed in metacercariae rather than sequencing or assembly limitations. Regardless, because stage 5 metacercariae are ready for transmission to the final host (Levri and Lively 1996), these data provide an excellent first step towards identifying the genes involved in this critical stage of parasite development.

**Table 1.** Assembly statistics for whole transcriptome assemblies and orthologous transcript assemblies for PA-*Microphallus* and PE-*Microphallus*.

| Assembly Statistic | Whole transcriptome |  | Ortholog transcriptome |  |
| --- | --- | --- | --- | --- |
|  | PA-*Microphallus* | PE-*Microphallus* | PA-*Microphallus* | PE-*Microphallus* |
| Number of transcripts | 15435 | 29564 | 7584 | 7584 |
| Minimum transcript length | 297 | 297 | 297 | 297 |
| Maximum transcript length | 10398 | 13584 | 9285 | 12381 |
| Median transcript length | 837 | 585 | 1173 | 1191 |
| Mean transcript length (+/- SD) | 1225.18 (1102.41) | 959.67 (1007.52) | 1505.35 (1157.72) | 1565.81 (1287.64) |
| Length of transcriptome (nt) | 18910587 | 28371675 | 11416539 | 11875112 |
| N content (%) | 0 | 0 | 0 | 0 |
| GC (%) | 47.25 | 48.05 | 47.22 | 47.21 |
| N50 | 1782 | 1368 | 2019 | 2070 |

We generated local BLAST databases for both annotated *Microphallus* transcriptomes that allow for simple queries of nucleotide and protein sequences. This resource will be broadly useful for future studies geared towards identifying candidate genes from *Microphallus* as well as other related ecologically and/or medically important parasites by providing an easy way to search for genes in a well-characterized host-parasite system. These databases will also be an asset for phylogenetic or molecular evolution-focused studies that would benefit from the addition of data from Trematoda.

### Ortholog identification and levels of genetic variation within and genetic differentiation between *Microphallus*isolates

We identified 7584 pairs of one-to-one orthologous transcripts between the two transcriptomes (Table S2) and 7851 and 21980 non-one-to-one orthologous transcripts for PA-*Microphallus* and PE-*Microphallus*, respectively. The two one-to-one ortholog transcriptomes contain higher mean transcript lengths (18.6% longer for PA-*Microphallus*; 38.7% longer for PE-*Microphallus*) and larger N50 values (11.7% greater N50 for PA-*Microphallus*; 33.9% greater N50 for PE-*Microphallus*) than their corresponding reference transcriptomes (Table S1). The longer transcripts and higher N50 values in the one-to-one ortholog sets relative to the reference transcriptomes indicate that the one-to-one orthologs represent a set of approximately full-length high-quality genes that occur once and only once in both transcriptome assemblies. More than half (PA-*Microphallus*: 51.7%; PE-*Microphallus*: 51.2%) of the BUSCO genes are retained in this reduced gene set, further indicating that our filtering steps likely removed poor-quality transcripts from our ortholog transcriptomes (Table 2). The annotation statistics for the ortholog transcriptomes are broadly similar to the annotation statistics for the reference transcriptomes (Table S1), suggesting the ortholog transcriptomes are likely representative of the reference transcriptomes.

**Table 2.** BUSCO assessment for transcriptome completeness. Includes 843 conserved metazoan genes. Of these 843 genes, we assessed the number of genes that were complete, duplicated, fragmented, and missing among our reference transcriptomes and ortholog transcriptomes.

|  | Whole transcriptome |  | Ortholog transcriptome |  |
| --- | --- | --- | --- | --- |
|  | *PA-Microphallus* | *PE-Microphallus* | *PA-Microphallus* | *PE-Microphallus* |
| # of complete single-copy genes (%) | 579 (68.7) | 573 (68.0) | 436 (51.7) | 432 (51.3) |
| # of duplicated genes (%) | 0 (0) | 2 (0.2) | 0 (0) | 0 (0) |
| # of fragmented genes (%) | 66 (7.8) | 122 (14.5) | 51 (6.1) | 55 (6.5) |
| # of missing genes (%) | 198 (23.5) | 146 (17.3) | 356 (42.2) | 356 (42.2) |

Of the 7584 transcripts in the ortholog transcriptomes, a distinct majority received both blastx and GO annotation for PA-*Microphallus* (5746 transcripts, 75.8%) and PE-*Microphallus* (5793 transcripts, 76.4%). An additional 15.9% (1211 transcripts) and 15.8% (1203 transcripts) of these transcripts received only blastx annotations for PA-*Microphallus* and PE-*Microphallus*, respectively. The species that were top blast hits most often and the species that were hit most frequently in blastx searches include other trematodes (five species) as well as non-trematode Platyhelminthes (two species) and other metazoans with well-annotated genomic and/or transcriptomic resources (Table S1). Species within Platyhelminthes were the top blastx hit for 90.6% (6337 transcripts) and 90.3% (6356 transcripts) of the orthologous transcripts that received blastx hits for PA-*Microphallus* and PE-*Microphallus*, respectively. The remaining 663 annotated orthologs for PA-*Microphallus* received top blast hits from mollusks (393 transcripts, 5.6%), other metazoans (253 transcripts, 3.6%), or non-metazoan taxa (17 transcripts, 0.2%), and the remaining 682 annotated PE-*Microphallus* transcripts received top blast hits from mollusks (418 transcripts, 5.9%), other metazoans (250 transcripts, 3.6%), or non-metazoan taxa (14 transcripts, 0.2%). The relatively small proportion of orthologous transcripts with a mollusk as their top hit could reflect low levels of contamination during the process of isolating metacercariae from snail tissue but is likely also linked in part to availability of well-annotated genomic and/or transcriptomic resources for the specific species that were hit (Table S1).

We used Popoolation to calculate nucleotide heterozygosity (Watterson’s θ) for each orthologous transcript by mapping PA-*Microphallus* and PE-*Microphallus* reads to their own ortholog transcriptomes and to each other’s ortholog transcriptomes, allowing us to account for differences that may result from the reference to which reads were mapped. Mean (+/− SD) θ per transcript for PA *Microphallus* mapped to the PA-*Microphallus* orthologs was 0.0044 (+/− 0.0039) and was 0.0038 (+/−0.0038) when mapped to the PE-*Microphallus* orthologs. Mean (+/− SD) per transcript for PE *Microphallus* mapped to the PE-*Microphallus* orthologs was 0.0051 (+/− 0.0046) and was 0.0051 (+/−0.0041) when mapped to the PA-*Microphallus* orthologs. These estimates of with observations from a previous allozyme-based estimate of are broadly consistent (0.001-0.021) in several populations of *P. antipodarum*-infecting *Microphallus* (Dybdahl and Lively 1996). All pairwise comparisons of medians (Mann-Whitney U-test) and distributions of values are significantly different between the two types of *Microphallus* (*p* < 0.001), regardless of which transcriptome reads are mapped. This result provides an initial line of evidence for higher genetic variation in PE-*Microphallus* relative to PA-*Microphallus*. For PE-*Microphallus*, the median (*p* = 0.898) and distribution (*p =* 0.578) of (Kolmogorov-Smirnov test) indicate that were not different from one another when mapped to either reference. By contrast, PA-*Microphallus* median (*p* < 0.001) and distribution (*p* < 0.001) of θ were significantly different when mapped to PA-*Microphallus vs*. PE-*Microphallus*, indicating thatIn order to account for the ortholog different SNPs may be called depending on the reference to which reads are mapped. One caveat to definitive interpretation of these results is the differences in sampling between the two types of *Microphallus* in our study: because the PA-*Microphallus* sample was produced from pooled RNA from multiple individuals, we cannot tie sequence differences back to individual worms. On the other hand, because we only sequenced a single PE-*Microphallus* individual, we can definitively ascribe within-sample variation to individual variation, but are not able to evaluate across-individual) and Popoolation2 (used to estimate *F*_*ST*_, below) are designed to handle pooled sequencing data and take into account the number of pooled individuals sequenced in the analyses.

We used Popoolation2 to calculate *F*_*ST*_ at each SNP between PA-*Microphallus* and PE-*Microphallus*. In order to account for the ortholog transcriptome to which reads were mapped, we calculated *F*_*ST*_ per-SNP between PA-*Microphallus* and PE-*Microphallus* mapped to the PA*Microphallus* ortholog transcriptome and calculated *F*_*ST*_ per-SNP between PA-*Microphallus* and PE*Microphallus* mapped to the PE-*Microphallus* ortholog transcriptome. We found a mean (+/− SD) *F*_*ST*_ per SNP between the two types of *Microphallus* mapped to the PA-*Microphallus* ortholog transcriptome of 0.129 (+/− 0.129) and a mean (+/− SD) *F*_*ST*_ per SNP between the two types of *Microphallus* mapped to the PE-*Microphallus* ortholog transcriptome of 0.148 (+/− 0.138). These *F*_*ST*_ values provide the first report of which we are aware of patterns of genetic differentiation between *P. antipodarum*-infecting and *P. estuarinus*-infecting *Microphallus*. Our *F*_*ST*_ estimates are an order of magnitude higher than the *F_ST_* values reported in an early allozyme-based study comparing *F*_*ST*_ across populations of *Microphallus* infecting *P. antipodarum* (Dybdahl and Lively 1996), which provides an initial line of evidence that there is marked genetic differentiation between these two *Microphallus* types relative to the across-population differentiation observed for *Microphallus* infecting *P. antipodarum.*

When mapping to the PA-*Microphallus* ortholog transcriptome, we identified 36 *F*_*ST*_ outlier-containing transcripts. Of these 36 transcripts, 31 received blastx and GO annotation, one received only blastx annotation, and four were unable to be annotated (Table S3). 31 of the annotated transcripts had top hit blastx hit species within Trematoda (Table S4). Based on Blast2GO annotations, these *F*_*ST*_ outlier-containing transcripts appear to be involved in cellular processes, metabolism, biological regulation, and localization (Table S5). When mapping to the PE-*Microphallus* ortholog transcriptome, we identified 30 *F_ST_* outlier-containing transcripts. Of these transcripts, 22 received both blastx and GO polymorphism. Despite these limitations, Popoolation (used to estimat annotations, three received only blastx annotation, and three were unable to be annotated (Table S3). All 25 of the annotated transcripts received top blastx hit species within Platyhelminthes, 24 of which were within Trematoda (Table S4). The annotated *F*_*ST*_ outlier-containing transcripts identified when mapping to the PE-*Microphallus* orthologs appear to be involved in similar biological processes as those identified by mapping to PA-*Microphallus* (Table S5).

Only two of the transcripts were identified as *F*_*ST*_ outliers in both comparisons. Both of these transcripts could represent genes that are evolving relatively quickly compared to their respective ortholog transcriptome and/or may be potential targets of selection in one or both *Microphallus* types. These candidate genes are N-alpha-acetyltransferase, which is involved in the N-terminal acetyltransferase C (NatC) complex and acetylation of amino acids, and permease 1 heavy chain, which is involved in maltose metabolic processes (Table S3). Genes involved in the NatC pathway have been implicated in modulating stress tolerance in *Caenorhabditis elegans* (Warnhoff and Kornfeld 2015). In schistosomes (Trematoda), permease 1 heavy chain is characterized as playing an important role in nutrient uptake from hosts and is expressed during all major schistosome life stages (Krautz-Peterson *et al.* 2007). This information from a related trematode group with a broadly similar life cycle indicates that permease 1 heavy chain might also play a role in nutrient acquisition in *Microphallus*. More directed study of patterns of molecular evolution and the roles that these two genes play in responses to physiological stresses associated with freshwater *vs.* estuarine environments (N-alpha-acetyltransferase) as well as nutrient uptake by parasites from hosts (permease 1 heavy chain) in *Microphallus* will provide more definitive insights into their functions.

### GO functional enrichment and KEGG pathway identification

We used Blast2GO to perform functional enrichment analyses comparing our ortholog transcriptomes against their associated reference transcriptomes, to identify over *vs.* underrepresented functional groups that are shared between the two types of *Microphallus*. We also used functional enrichment analyses of our non-one-to-one orthologs against their associated reference transcriptomes to assess the types of genes that are over *vs.* underrepresented among the transcripts that did not represent one-to-one orthologs between PA-*Microphallus* and PE-*Microphallus.* We identified seven GO terms that were significantly underrepresented among orthologous transcripts in PA-*Microphallus* relative to the PA-*Microphallus* reference transcriptome. These same seven functional groups are significantly overrepresented among the non-one-to-one orthologs for PA-*Microphallus* relative to the PA-*Microphallus* reference transcriptome (Table S6). For the PE-*Microphallus* orthologs relative to the PE-*Microphallus* reference, we identified 20 significantly underrepresented and six significantly overrepresented functional groups. There were five underrepresented and 14 overrepresented functional groups among the non-one-to-one PE-*Microphallus* orthologs relative to the PE-*Microphallus* transcriptome. All but four of the functional groups among the PE-*Microphallus* transcripts were only significantly over or underrepresented among either the one-to-one or non-one-to-one orthologs but were never significantly over/underrepresented in both transcript sets (Table S6).

There were only two GO categories that were that were over or underrepresented in both types of *Microphallus*. First, genes with functions related to hydrolase activity are underrepresented among one-to-one orthologs and overrepresented among non-one-to-one orthologs for both PA-*Microphallus* and PE-*Microphallus*. The presence of genes involved in hydrolase activity among genes that were significantly functionally enriched is interesting in light of evidence that hydrolases are involved in damaging epithelial lining of the digestive tracts of hosts by the parasitic nematode *Contracaeum rudolphii* (Dziekonska-Rynko and Rokicki 2005). Second, genes involved in carbohydrate metabolic processes are underrepresented among one-to-one orthologs for both types of *Microphallus* relative to the reference transcriptomes. Evidence that schistosome parasites dynamically shift carbohydrate metabolic processes throughout the course of their life cycle (Horemans *et al.* 1992) hint that these genes could also be of relevance to *Microphallus*. Altogether, both of these functional categories of genes provide a promising set of candidate genes for assessment of the changes in gene expression and physiological responses associated with the infection process in *Microphallus*. The marked differences in functional groups that are significantly over *vs.* underrepresented in the ortholog transcriptomes relative to the reference transcriptomes between the two *Microphallus* types suggests that the types of genes present in the two reference transcriptomes are often different. Whether this result is more a consequence of biological differences between PA-*Microphallus* and PE-*Microphallus* or differences in sequencing coverage cannot be determined at this time but warrants future study.

Finally, we performed KEGG analyses as implemented in Blast2GO to identify the types of enzyme pathways that were being expressed in each transcriptome. This analysis identified 123 KEGG pathways in the PA-*Microphallus* transcriptome and 128 KEGG pathways in the PE-*Microphallus* transcriptome (Table S7). All but 11 of the KEGG pathways that we identified were expressed in both transcriptomes. Most of the 122 pathways present in both *Microphallus* transcriptomes are involved in various metabolic processes (49 pathways, 40.2%) or biosynthetic processes (41 pathways, 33.6%). Of the 11 pathways only identified in a single transcriptome, the three pathways identified in only PA-*Microphallus* include the riboflavin metabolic pathway, the penicillin and cephalosporin biosynthesis pathway, and the biosynthesis of siderophore group nonribosomal peptides. The eight pathways that were identified in only PE-*Microphallus* include a variety of biosynthesis and metabolic processes (Table S7). One such pathway, glycosphingolipid biosynthesis, has been implicated in response to salinity stress in *Litopenaeus vannamei*, the Pacific white shrimp (Chen *et al.* 2015). This result is suggestive of a scenario where these genes contribute to the ability of PE-*Microphallus* to tolerate salinity fluctuations associated with estuarine habitats. This important habitat difference between PA-*Microphallus* and PE-*Microphallus* is likely to differentially influence the selective pressures experienced by these two parasites and thus has the potential to influence patterns and rates of molecular evolution (*e.g.*, Mitterboeck *et al*. 2016), making genes involved in salinity tolerance and/or stress particularly interesting candidates for future study.

### Predictions of microsatellite loci and primers

We used MISA software to identify SSRs within the ortholog transcriptomes and used Primer3 and BLAST to predict primers that could be used for future microsatellite-based analyses of *Microphallus*. We identified 109 SSRs in 100 transcripts for PA-*Microphallus* and 117 SSRs in 107 transcripts for PE-*Microphallus*. We identified all six types of microsatellites for which we searched, with trinucleotide sequences with at least five repeats representing the most common microsatellite type (62.3% of SSRs identified in PA-*Microphallus* and 60.7% of SSRs identified in PE-*Microphallus*) (Table 3; Fig. S1). Eight and four SSRs, respectively, were defined as compound for PA-*Microphallus* and PE-*Microphallus* (Table 3). We also used perl scripts available with MISA (http://pgrc.ipkgatersleben.de/misa/) to compile specific details about the primer sets (*e.g.*, location, size of product, melting temperature). We generated predicted primers for 84 of the PA-*Microphallus* SSRs from 100 sequences and for 83 of the PE-*Microphallus* SSRs from 113 sequences (Tables S8, S9).

**Table 3.**
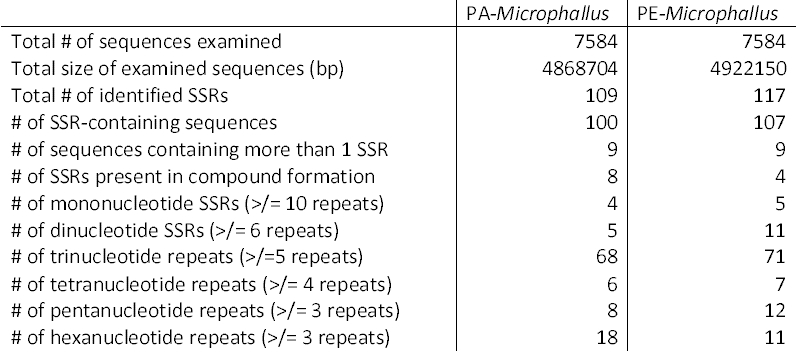
Summary of potential microsatellite loci for the PA-*Microphallus* and PE-*Microphallus* one-to-one ortholog transcriptome assemblies.

|  | PA-*Microphallus* | PE-*Microphallus* |
| --- | --- | --- |
| Total # of sequences examined | 7584 | 7584 |
| Total size of examined sequences (bp) | 4868704 | 4922150 |
| Total # of identified SSRs | 109 | 117 |
| # of SSR-containing sequences | 100 | 107 |
| # of sequences containing more than 1 SSR | 9 | 9 |
| # of SSRs present in compound formation | 8 | 4 |
| # of mononucleotide SSRs ($\geq 10$ repeats) | 4 | 5 |
| # of dinucleotide SSRs ($\geq 6$ repeats) | 5 | 11 |
| # of trinucleotide repeats ($\geq 5$ repeats) | 68 | 71 |
| # of tetranucleotide repeats ($\geq 4$ repeats) | 6 | 7 |
| # of pentanucleotide repeats ($\geq 3$ repeats) | 8 | 12 |
| # of hexanucleotide repeats ($\geq 3$ repeats) | 18 | 11 |

Future studies will be able to use these markers to characterize, for example, population genetic structure within and genetic differentiation between these two types of *Microphallus* isolated from two ecologically distinct snail taxa. Comparing patterns of molecular evolution among these orthologous genes from more *Microphallus* individuals isolated from *P. antipodarum* and *P. estuarinus* will allow for research directed at characterizing the strength and efficacy of selection and the importance of population structure in the context of a prominent model for host-parasite interactions and coevolution. Researchers can also use these resources to perform cost-effective analyses incorporating many *Microphallus* individuals from multiple populations to address key questions regarding the evolutionary connections between these two types of *Microphallus* (*e.g.,* whether there is gene flow between the *Microphallus* infecting *P. antipodarum* and *P. estuarinus* and whether there is similar evidence for local adaptation and coevolution between *P. estuarinus* and *Microphallus* as exists for *P. antipodarum* and *Microphallus*).

## Conclusions

We provide a set of widely useful transcriptomic and genomic resources for an evolutionarily and ecologically important group of parasites. *Potamopyrgus antipodarum* and *Microphallus livelyi* are a textbook system for the study of host-parasite coevolutionary interactions, and the data generated herein represent the first large-scale genome-based resources for *Microphallus*. The annotated transcriptome assemblies, orthologous gene sets, blast databases, SNPs, and microsatellite loci and primers will facilitate research for scientists interested in host-parasite coevolution, local adaption, and/or patterns of molecular evolution and phylogenetics of these and other trematode parasites. Our *F_ST_*, functional enrichment, and KEGG pathway analyses provide a promising set of candidate genes and pathways for future study and further characterization for potential roles in physiological and stress responses relevant to organisms inhabiting freshwater *vs.* estuarine environments, as well as genes involved in parasitic infection of host taxa.

## Acknowledgements

We thank Gery Hehman for RNA sequencing assistance, Curt Lively, Daniela Vergara, Katelyn Larkin, and Mike Winterbourn for field collections, and Joel Sharbrough for the custom python script (github.com/jsharbrough/grabContigs). This project was funded by Sigma-Xi (grant number 18911100 BR01; L. Bankers), the Iowa Science Foundation (grant number 14-03; M. Neiman and L. Bankers), the National Geographic Society (grant number 9595-14; K. Larkin) and the National Science Foundation (NSF-MCB grant number 1122176; M. Neiman).

